# G2M splits mouse embryonic stem cells into naïve and formative pluripotency states

**DOI:** 10.1101/616516

**Authors:** Kersti Jääger, Daniel Simpson, Maria Kalantzaki, Angela Salzano, Ian Chambers, Tamir Chandra

**Affiliations:** MRC Centre for Regenerative Medicine, Institute for Stem cell Research, School of Biological Sciences, University of Edinburgh, Edinburgh EH16 4UU, UK; MRC Institute of Genetics and Molecular Medicine, Western General Hospital, The University of Edinburgh, Edinburgh, EH4 2XU, UK

## Abstract

Embryonic stem cells (ESCs) express heterogeneous levels of pluripotency and developmental transcription factors (TFs) and their cell cycle is unsynchronised when grown in the presence of serum. Here, we asked whether the cell cycle and developmental heterogeneities of ESCs are coordinated by determining the state identities of G1- and G2M-enriched mouse ESCs (mESCs) at single cell resolution. We found that G2M cells were not all the same and demonstrate their split into the naïve and formative (intermediate) pluripotency states marked by high or low Esrrb expression, respectively. The naïve G2M sub-state resembles ‘ground’ state pluripotency of the LIF/2i cultured mESCs. The naïve and formative G2M sub-states exist in the pre- and post-implantation stages of the mouse embryo, respectively, verifying developmental distinction. Moreover, the G2M sub-states partially match between the mouse and human ESCs, suggesting higher similarity of transcriptional control between these species in G2M. Our findings propose a model whereby G2M separates mESCs into naïve and formative pluripotency states. This concept of G2M-diverted pluripotency states provides new framework for understanding the mechanisms of pluripotency maintenance and lineage specification in vitro and in vivo, and the development of more efficient and clinically relevant reprogramming strategies.

## INTRODUCTION

### Heterogeneity of mESC populations and pluripotency state transitions

Different pluripotency states present in the peri-implantation mouse embryo are recapitulated *in vitro* in the form of mouse embryonic stem cells (mESCs, E4.5, naïve state), epiblast-like cells (EpiLCs, E5.5, early post-implantation or formative state) and epiblast stem cells (EpiSCs, E6.5, primed state)[1]. The regulatory networks controlling the transitions between these states include combinatorial and dosage-dependent interactions between general and state-specific pluripotency and priming transcription factors (TFs). The expression of pluripotency TFs including Nanog [2], Esrrb [3], Zfp42/Rex1 [4] is not uniform in mESCs when grown in the presence of serum and leukaemia inhibitory factor (LIF) [5]. Therefore, according to the emerging model, the naïve and primed cell states, as well as the recently described intermediate or formative state, co-exist and are interconvertible in these culture conditions. How these transitions occur in individual cells and whether they follow stochastic or deterministic regulatory principles, is still under active investigation.

### Cell cycle and fate choice

Cell cycle phasing has been shown to contribute to the observed heterogeneity of mESCs, and efforts have been made to separate cell cycle driven heterogeneity from construction of TF circuitries of different developmental states [6,7]. These approaches have intuitively assumed that the ESC transcriptomes within the same cell cycle phase are more uniform than between phases. Given the dynamic nature of pluripotency state transitions, pluripotency and priming TFs could be variably expressed in the same cycle phase. Indeed, several studies have dissected the role of a particular cell cycle stage in determining ESC fate choice, and consequently shown that ESCs can adopt different fates when positioned in a certain cell cycle phase [8–11]. These findings imply potential diversion and switching between distinct gene regulatory programs in the same cell cycle phase.

Here, we asked whether mESCs residing in a particular cell cycle phase are in the same transcriptional state space or diverge into distinguishable naïve and primed sub-states (**Fig. 1A**) that facilitate change in cell fate. We focussed our attention on G1 and G2M phases that have been shown to either facilitate cell differentiation [10] or enhance maintenance of pluripotency [8], respectively. To our knowledge, this is the first attempt to dissect the cell cycle-centred TF heterogeneity of ESCs.

**Fig. 1.**
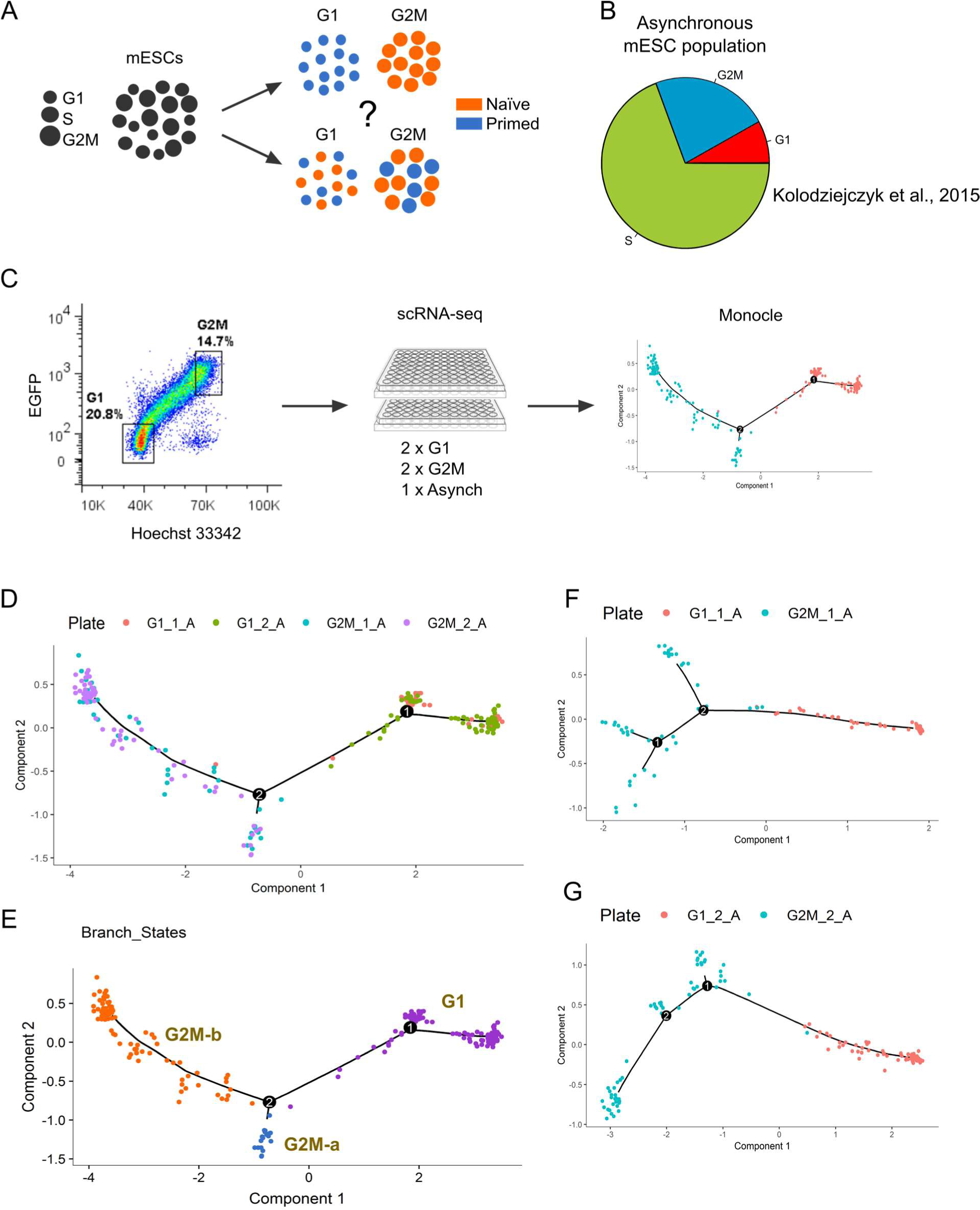
G2M splits mESCs into sub-states. **A**. We asked whether serum/LIF-grown mESCs diverge into distinct pluripotency sub-states when positioned in either G1 or G2M cell cycle phase. **B**. Cyclone-predicted cell cycle distribution of an asynchronous mESC population from Kolodziejczyk et al., 2015 showing mESC-characteristic cell cycle profile with the majority of cells in S phase. **C.** Single cells were sorted by flow cytometry into 96-well plates using the indicated G1 and G2M gates (or asynchronous cells) and applied to single-cell RNA sequencing followed by branching trajectory analysis using Monocle. **D.** G1- and G2M enriched cells from two plates (G1 – red/green; G2M – blue/purple) were ordered by pseudotime on a branching trajectory revealing two nodes where either G1 (node 1) or G2M (node 2) cells split into two sub-populations. **E.** Same clustering as in D using color-coding for G1 (purple), G2M-a (blue) and G2M-b (orange) states. **F** and **G**. Branching trajectory constructed for replicate plates (plate 1 in F and plate 2 in G) showing bifurcation of G2M cells on both assay plates.

## EXPERIMENTAL PROCEDURES

### mESC culture

mESCs were cultured on 0.1% gelatin-coated tissue culture plastic in GMEM/β-mercaptoethanol/10%FCS/LIF as described before [12].

### Generation of mESCs expressing endogeneous cell cycle reporter

For the generation of endogenous Cyclin B1-GFP reporter cell line, a targeting strategy similar to Bressan et al., 2017 was used [13]. 1 kb of homologous sequences upstream and downstream of *CCNB1* stop codon were amplified from wild type (wt) E14Tg2A ESCs. A targeting cassette with fluorescent reporter was generated using Gibson assembly in the following configuration: 5’-1 kb homologous sequence – flexible linker – enhanced green fluorescent protein (EGFP) – STOP codon – 1 kb of 3’ homologous flanking region. Independent plasmids coding for CRISPR/Cas9-Nickase (D10A) and sgRNAs (CTGCAGATGTAGCAGTCTAT, ACCATGTGCCGCCTGTACAT) were used for introducing an EGFP tag to the C-terminus of *CCNB1* in E14Tg2A ESCs. For the delivery, 3×10^6^ mESCs were seeded onto a 10-cm dish and transfected using Lipofectamine 3000 (Invitrogen) according to manufacturer’s instructions. Cells were transfected with 2 ug of Cas9-Nickase (D10A) expression vector, 1 ug of each gRNA expression vector, and 700 ng of donor cassette. The representation of the targeted *CCNB1* locus is shown in **Fig. S1A**. The efficiency of genome editing was analysed by flow cytometry 48h post-transfection with 0.2% of the total population being positive for GFP expression. It is important to note that this percentage only describes targeted cells that were in S/G2/M phase at the time of analysis due to natural Cyclin B1 expression dynamics during cell cycle. GFP-positive cells were sorted as single cells into 1x 96-well plate and clones expanded for 10 days. Cells with ESC like morphology were picked and 7 clones successfully expanded for genomic DNA isolation using DNeasy Blood and Tissue Kit (Qiagen) followed by genotyping using Taq DNA polymerase (Takara) according to manufacturer’s instructions. Primers used for genotyping were: Primer 1: 5’-CCTAATATGGAAGGAGTGTGAC-3’, Primer 2: 5’-AGGAATACTGTTGATAAAATG-3’. Out of 7 analysed clones, one was homozygous (N7) and one heterozygous for *CCNB1* targeting (**Fig. S1B**). Clone N7 and one wt clone (N5) were selected for FACS analysis (**Fig. S1C**). Snapshots taken form a time-lapse imaging of N7-mESCs confirmed cell cycle regulated expression dynamics of CyclinB1-GFP fusion protein (**Fig. S1D**).

### Cell sorting and single-cell RNA-seq library preparation

N7-mESCs were stained with 50 uL Hoechst 33342 (50 mg/mL stock) in 10 mL medium 45 min at 37°C prior to FACS-sorting of single cells directly into 2 uL of lysis buffer (0,2 % Triton X 100, 2U/uL RNase inhibitor in water) in 96-well plates. Plates were immediately put on dry ice and stored at −80°C until library preparation. 2x 96-well plates were sorted from both G1 and G2M gate (**Fig. 1C**), and 1x 96-well plate form non-gated asynchronous population. We used Smart-seq2 protocol for cDNA synthesis [14], and Fluidigm protocol (PN 100-7168 M1) for tagmentation and index-PCR using Nextera chemistry to prepare single cell sequencing libraries. 2x 96-plates were pooled together and sequenced on one lane on Illumina HiSeq 4000 (50 bp, single-end reads).

### Read processing and quality control

Raw reads were mapped against the Ensembl mouse reference genome version GRCm38.68 using STAR RNA-seq aligner [15] with the inclusion of the sequences for the External RNA Controls Consortium (ERCC) spike-ins. Mapped reads per gene were calculated using *Rsubread* [16]. Genes with zero counts across all cells were excluded. Standard quality control metrics were calculated using the *scater* package [17]. The following quality control criteria was used: cells with more than 25% of reads mapping to ERCCs or fewer than 100,000 total reads or fewer than 4,000 genes detected (at least one read per gene) were excluded from downstream analyses (**Fig. S2A-D**). The number of cells passing this quality control were: 62 cells for Asynch, 111 cells for G1 and 104 cells for G2M.

### Statistical analyses

The input genes for the construction of the G1-G2M pseudotime trajectory were selected by differential expression between the G1 and G2M cells using the *differentialGeneTest* function in Monocle 2 [18]. Log2 +1 counts per million (CPM) values were computed using the *calculateCPM* function in *scater*. Boxplots were constructed using *ggplot2* and statistical tests applied using *ggpubr*. A Wilcox statistical test was applied in pairwise comparisons and a Kruskal-Wallis test in group comparisons. Differential expression analysis was conducted using the *SCDE* package in R [19].The resulting z-score ranked gene lists were used in a GSEA analysis [20].

### Comparisons between different datasets

Raw counts from the [6] were downloaded from http://www.ebi.ac.uk/teichmann-srv/espresso/index.php, and cells with more than 25% of reads mapping to ERCCs or fewer than 10^6.3^ reads or fewer than 9,000 genes detected (at least one read per gene) were rejected, resulting in 668 cells. The total number of cells went down from the original of 704 cells to 668 primarily due to the new ERCC threshold set. Data from [21] was obtained from the Gene Expression Omnibus (GEO) database under a reference number GSE100597, and the associated metadata was obtained using the *getGEO* function from the *GEOquery* package. No further quality criteria were applied. Data from [22] was downloaded from EMBL-EBI ArrayExpress under the accession number E-MTAB-6819. Human gene names were converted to mouse names using BioMart. *scmapCluster()* from the *scMap* package was used for all comparisons between our and external datasets [23].

## RESULTS

### G2M splits mESCs into sub-states

To begin to elucidate the heterogeneity of cell cycle phase-matching ESCs, we aimed to incorporate into our analyses a comparable number of cells from both G1 and G2M. Cell cycle phase assignment using Cyclone [24] of an asynchronous population of mESCs [6] revealed the majority of cells to reside in S phase (**Fig. 1B**), a cell cycle profile characteristic to ESCs [25]. Therefore, to capture mESCs in G1 and G2M phases, we generated a mESC line that expressed GFP-tagged Cyclin B1 from its endogeneous loci (**Fig. S1A**). The expression of Cyclin B1 protein directly correlates with cell cycle progression by peaking in G2M and rapidly disappearing in G1, and when combined with a fluorescent tag and Hoechst 33342 live DNA staining, provides a simple single-reporter system for separating G1 and G2M from S phase using flow cytometry (**Fig. 1C, Fig. S1D**). This approach enabled us determine gene expression in chemically unperturbed cell cycle enriched ESC populations.

Sorted cells were applied to single cell sequencing library preparation using a modified version of the Smart-seq2 protocol by Picelli et al., 2014 (see **Experimental procedures**). We obtained 111 cells in G1 and 104 cells in G2M after quality filtering of cells collected from duplicate plates. We also sequenced one plate corresponding to the asynchronous population (62 cells). We used Cyclone to quantitatively assess the G1 and G2M enrichment in sorted ESCs (**Fig. S3A-C**). G1 cells appeared less uniform in terms of G1-signature (**Fig. S3B**), whereas 96 % of G2M-sorted ESCs were assigned to be in G2M and as little as 1 % or 3 % in G1 or S, respectively, confirming high purity of the sorted G2M fraction (**Fig. S3C**).

Next, to reconstruct cell-to-cell relationship of G1 and G2M-enriched mESCs on a branching trajectory, we used Monocle that orders individual cells by pseudotime and thereby enables tracking the emergence of hidden developmental states [18]. Curiously, we identified two branching points upon joint clustering of G1- and G2M-enriched cells (**Fig. 1D and E**), and the emergence of two G2M sub-states – G2M-a and G2M-b. To confirm plate-independent branching, we performed separate clustering of replicate G1 and G2M plates and saw a) continuous spreading of G1 cells along the trajectory, and b) bifurcation happening only in G2M on both plates (**Fig. 1F and G**). This finding clearly shows that not all cells in the G2M are the same.

### G2M sub-states separate by reciprocal expression of Esrrb and Pou3f1

To investigate the split of G2M in more detail, we performed unbiased single-cell consensus clustering of G2M cells using SC3 [26] which recovered near-perfect separation of G2M-a and G2M-b sub-states into two clusters (**Fig. 2A**). We identified 330 marker genes for the clusters (**Suppl. Table 1**) including 134 high-confidence genes in cluster 1 (q < 0.05) and 18 genes in cluster 2 (q < 0.05). Among the marker genes were the early post-implantation genes Pou3f1 and Fgf5 in cluster I, and naïve pluripotency genes Esrrb and Klf2 in cluster II (**Fig. 2A**). Opposite expression of early post-implantation and naïve genes has been shown to distinguish pre- and post-implantation epiblast of the mouse embryo and their corresponding cellular derivates *in vitro* [27,28] suggesting separation of G2M sub-states by different developmental state.

**Fig. 2.**
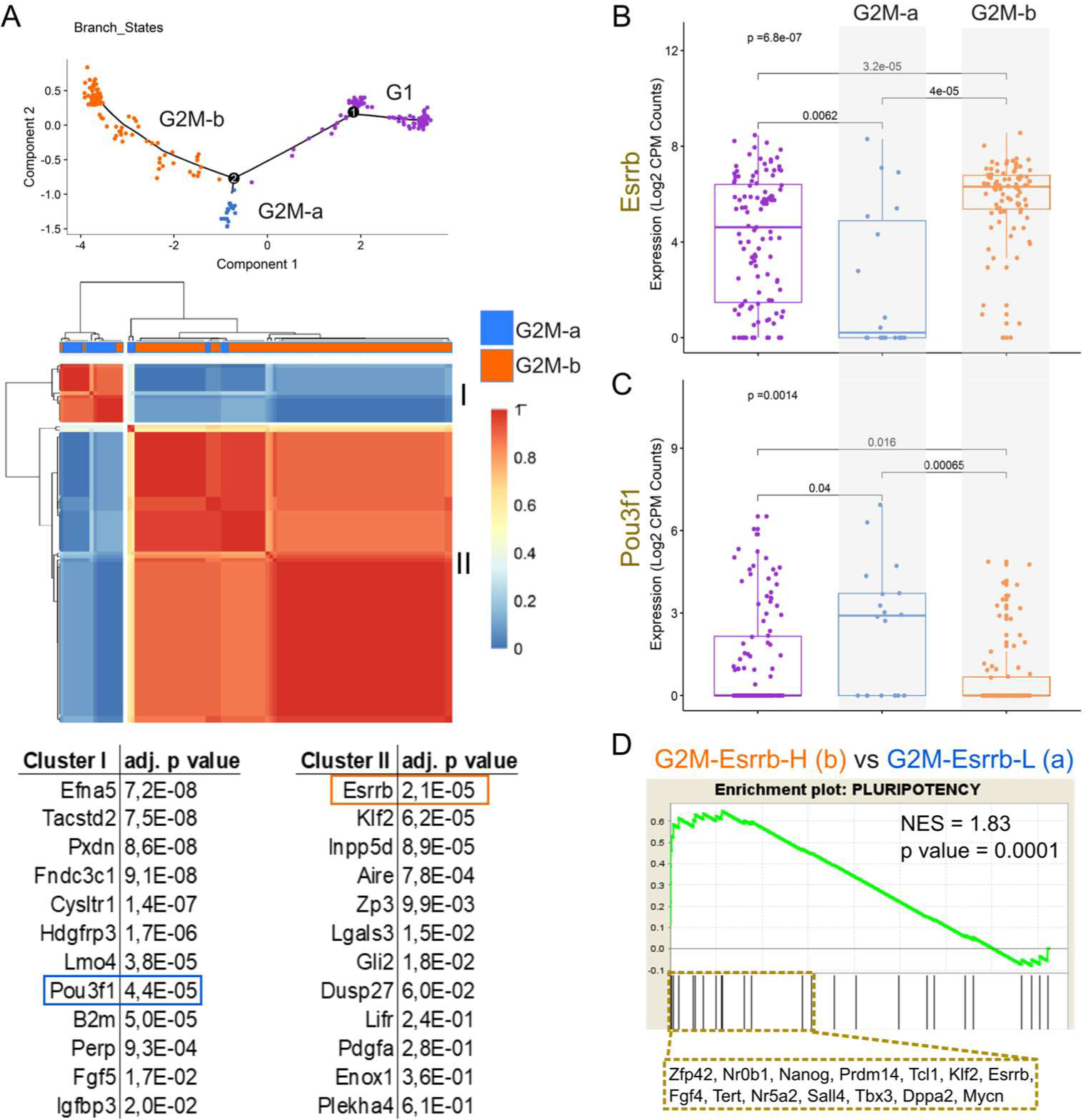
G2M sub-states separate by reciprocal expression of Esrrb and Pou3f1. **A.** Single-cell consensus heatmap showing separate clustering of G2M-a and G2M-b sub-states based on whole transcriptome expression. Selected significant marker genes in clusters I and II are shown below. The top gene Esrrb in cluster II is highlighted. **B** and **C.** Distribution of Esrrb (B) and Pou3f1 (C) expression in G1, G2M-a and G2M-b sub-states showing significantly higher expression of Esrrb in the G2M-b sub-state compared with G2M-a. **D.** Gene set enrichment analysis (GSEA) of pluripotency genes between G2M-Esrrb-high (H) and G2M-Esrrb-low (L) sub-states, showing strong and significant (p value = 0.0001) enrichment in the G2M-Esrrb-H. The list of the ‘leading edge’ genes is shown below. NES = normalised enrichment score.

The expression of Esrrb was distributed differently across individual cells at distinct Monocle states, confirming lower median expression (log2Ex < 1) in G2M-a and higher median expression (log2Ex > 6) in G2M-b (**Fig. 2B**). Reverse distribution of Pou3f1 in G2M-a (median log2Ex > 2) and G2M-b (median log2Ex = 0) confirmed reciprocal expression of Esrrb and Pou3f1 in G2M sub-states (**Fig. 2C**).

We conducted single-cell differential expression analysis, SCDE [19] between G2M-b (G2M-Esrrb-<u>H</u>igh) and G2M-a (G2M-Esrrb-<u>L</u>ow) and used the resulting ranked gene list (**Suppl. Table 2**) in a gene set enrichment analysis (GSEA) [20,29] to verify that the Esrrb expression positively correlated with the expression of the naïve pluripotency genes (**Fig. 2D**). These results indicate towards G2M-Esrrb-L and G2M-Esrrb-H split into alternative states of pluripotency.

### G2M sub-states represent features of the naïve and formative pluripotency states

To compare more directly whether the G2M-Esrrb-L and G2M-Esrrb-H differed by expression signatures of known pluripotency states (naïve, early post-implantation or formative, primed), we plotted the z-scores from the SCDE analysis for selected gene sets combined from [21,28,30,31] (**Fig. 3A**). To our excitement, a clear distinction between G2M-Esrrb-L and G2M-Esrrb-H appeared only by the expression of naïve and early post-implantation or formative state genes, where G2M-Esrrb-H cells expressed higher levels of Zfp42, Nanog, Prdm14 among others (positive z-scores) and G2M-Esrrb-L cells higher levels of Pou3f1, Fgf5, and Perp (negative z-scores, **Fig. 3A**). Such clear diversion was not seen between G1 sub-states (**Fig. S3D**). Importantly, both G2M-Esrrb-L and G2M-Esrrb-H cells expressed similar levels of core pluripotency TFs including Pou5f1 and Sox2 revealing general pluripotency in both G2M sub-states. Importantly, the expression of Otx2, Sox3 and several other priming genes was similar or slightly higher in the naïve G2M-Esrrb-H state than in the formative G2M-Esrrb-L state, suggesting identification of a very early exit-from-naivety state in G2M.

**Fig. 3.**
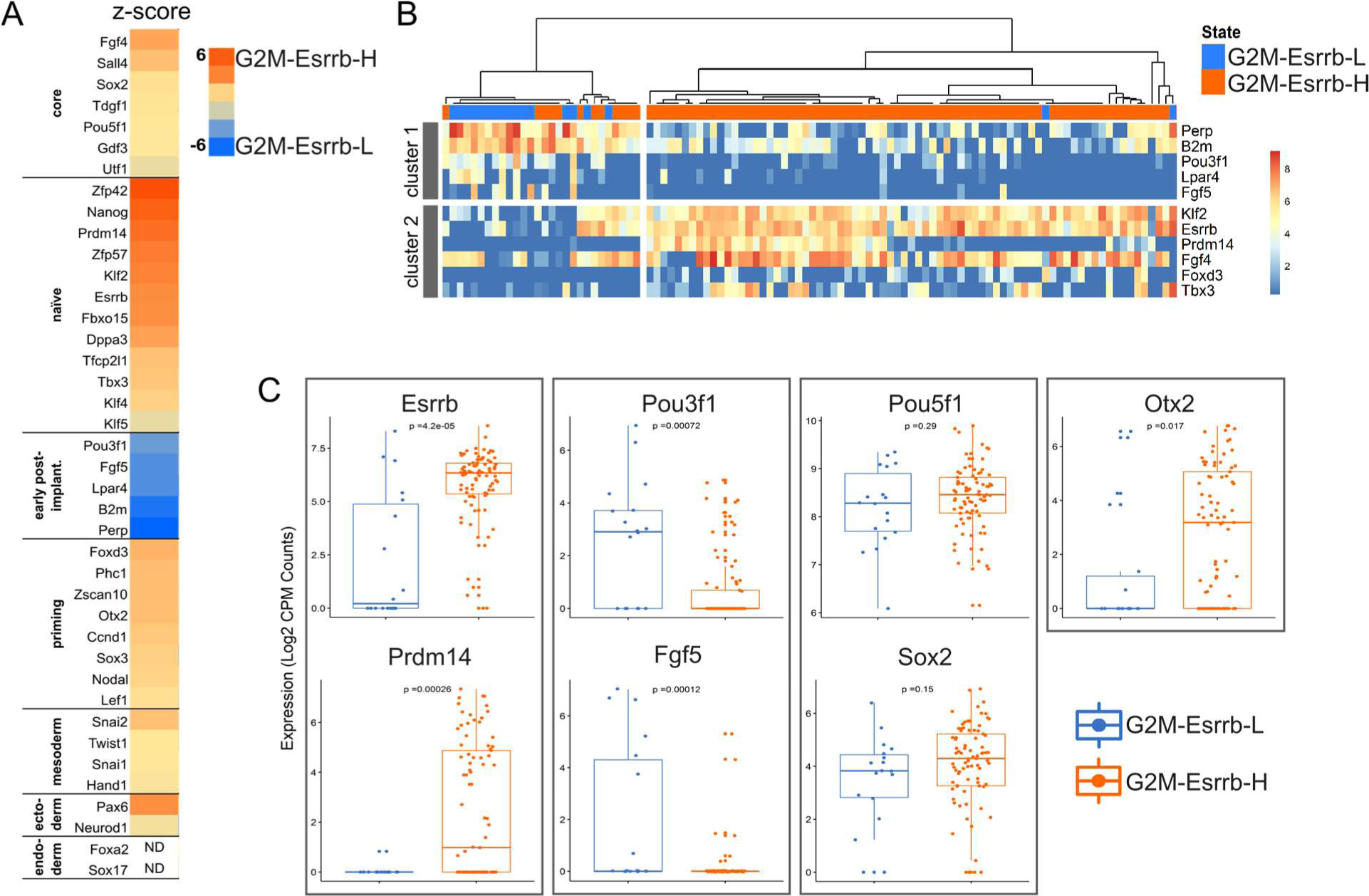
G2M sub-states represent features of the naïve and formative pluripotency states. **A.** Single-cell differential expression (SCDE) analysis between G2M-Esrrb-H and G2M-Esrrb-L sub-states showing opposite expression of naïve and early post-implantation genes, and similar expression of core pluripotency and priming genes. Higher z-score depicts higher expression in G2M-Esrrb-H compared to G2M-Esrrb-L, and vice versa. **B.** Single-cell clustered heatmap based on genes in 3A, showing higher expression of early post-implantation (formative pluripotency) marker genes (including Pou3f1 and Fgf5) in cluster 1, and higher expression of naïve pluripotency marker genes (including Esrrb and Klf2) in cluster 2. Other genes in 3A were not identified as significant marker genes of the two sub-states. **C.** Boxplots showing the distribution of expression of indicated genes across G2M-Esrrb-H and G2M-Esrrb-L populations in individual cells.

We visualised the expression of the genes from Fig3A on a clustered SC3 expression matrix which verified separation of G2M-Esrrb-L and G2M-Esrrb-H states solely by the reciprocal expression of naïve (cluster 2) and formative genes (cluster 1, **Fig. 3B**). This was confirmed by the distribution of expression of selected marker genes across G2M-Esrrb-L and G2M-Esrrb-H populations (**Fig. 3C**).

### mESC separation into G2M-Esrrb-H and G2M-Esrrb-L states is common in serum/LIF cultures

To test whether the split into G2M-Esrrb-H and G2M-Esrrb-L states is a general phenomenon characteristic to mESC populations cultured in serum/LIF, we re-analysed another dataset from asynchronous mESC population [6] marked by the name “Kolo”. We selected G2M-classified mESCs (using Cyclone) grown in serum/LIF (37 cells) from the Kolo dataset for analysis.

To enable direct matching of cells from two datasets, we used scMap [23], a tool that can project cells from one experiment onto the sub-types identified in another experiment. When mapping G2M-serum/LIF Kolo mESCs onto our identified cell cycle sub-states (including both G1 and G2M sub-states), we identified a match with both G2M-Esrrb-H (18 cells) and G2M-Esrrb-L (12 cells) states, verifying the G2M bifurcation (**Fig. 4A**). Seven cells out of the 37 Kolo G2M-serum/LIF cells were not assigned to any of our sub-states (unassigned, **Fig. 4A**).

**Fig. 4.**
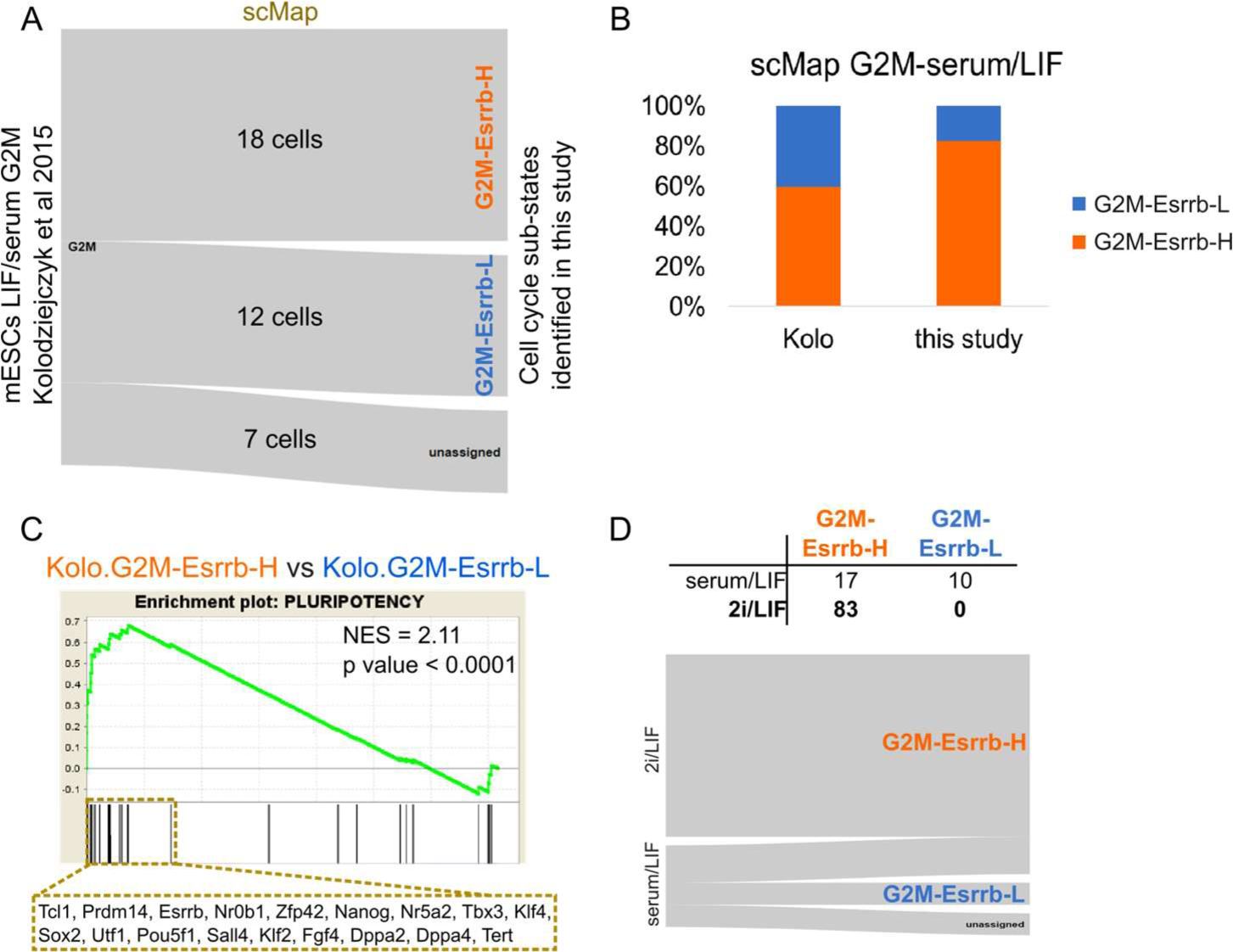
mESC separation into G2M-Esrrb-H (naïve) and G2M-Esrrb-L (formative) states is common in serum/LIF conditions. **A.** mESCs grown in serum/LIF from Kolodziejczyk et al 2015 (Kolo, G2M-classified) show mapping onto G2M-Esrrb-H (18 cells) and G2M-Esrrb-L (12 cells) sub-states. 7 of the Kolo G2M cells did not match any of our cell cycle states (unassigned). **B.** The proportions of G2M-Esrrb-H and G2M-Esrrb-L cells in the Kolo and our analysis of serum/LIF grown mESCs. **C**. GSEA of pluripotency genes between G2M-Esrrb-H and G2M-Esrrb-L sub-states in the Kolo dataset, showing strong and significant (p value < 0.0001) enrichment in the G2M-Esrrb-H. The list of the ‘leading edge’ genes is shown below. **D.** The number of G2M cells from the Kolo analysis grown either in serum/LIF or 2i/LIF that map to G2M-Esrrb-H (17 and 83 cells, respectively) and G2M-Esrrb-L (10 and 0 cells, respectively) sub-states. Note that all mESCs grown in 2i/LIF mapped to the G2M-Esrrb-H sub-state, suggesting G2M-Esrrb-H to resemble ’ground state’ pluripotency. NES = normalised enrichment score.

To compare the distributions into G2M-Esrrb-H and G2M-Esrrb-L sub-states between the Kolo and our data, we plotted the percentages of cells in each state on a stacked barplot (**Fig. 4B**). Slightly higher proportion of G2M cells were at the G2M-Esrrb-H state in our dataset (83 %) than in the Kolo dataset (60 %), possibly reflecting differences in cell cycle sorting versus cell cycle prediction, respectively, or in the duration of G2M. We performed GSEA between the Kolo G2M-Esrrb-H and G2M-Esrrb-L cells and observed similarly strong enrichment of the pluripotency gene expression in the G2M-Esrrb-H compared with G2M-Esrrb-L (**Fig. 4C**).

It has been demonstrated that mESCs grown in the presence of GSK-3α/β and MEK1/2 inhibitors (“2i”) mimic “ground-state” pluripotency, a state resembling conditions of the pre-implantation embryo [35]. To test whether the naïve G2M-Esrrb-H state is more similar to the ground state pluripotency, we performed mapping of the Kolo mESCs cultured both in 2i/LIF and serum/LIF onto our G2M sub-states. Indeed, all analysed Kolo 2i/LIF cells (G2M-classified) matched with the G2M-Esrrb-H state (83 cells), whereas the Kolo serum/LIF cells matched either with G2M-Esrrb-H (17 cells) or G2M-Esrrb-L (10 cells), or were unassigned (10 cells, **Fig. 4D**). This result shows that the ‘ground state’ of pluripotency emerges in serum/LIF cultured mESCs when in the early G2M phase.

### G2M sub-states separate the earliest pre-implantation stages

The naïve and formative G2M sub-states appear to match with the embryonic stages of the mouse development in vivo and best distinguish the earliest E3.5 and E4.5 developmental stages as evidenced by mapping of the data from Mohammed et al., 2017 [21] onto our identified pluripotency sub-states (**Fig. 5A**). Moreover, despite distinct growth requirements of the mouse and human naïve ESCs [36,37], we saw relatively strong matching between G2M-ESCs from the two species (**Fig. 5B**) by reverse mapping of our G2M sub-states onto G2M-classified hESCs [22], suggesting more conserved regulatory mechanisms of pluripotency maintenance in G2M that could remain unnoticed when analysing bulk populations.

**Fig. 5.**
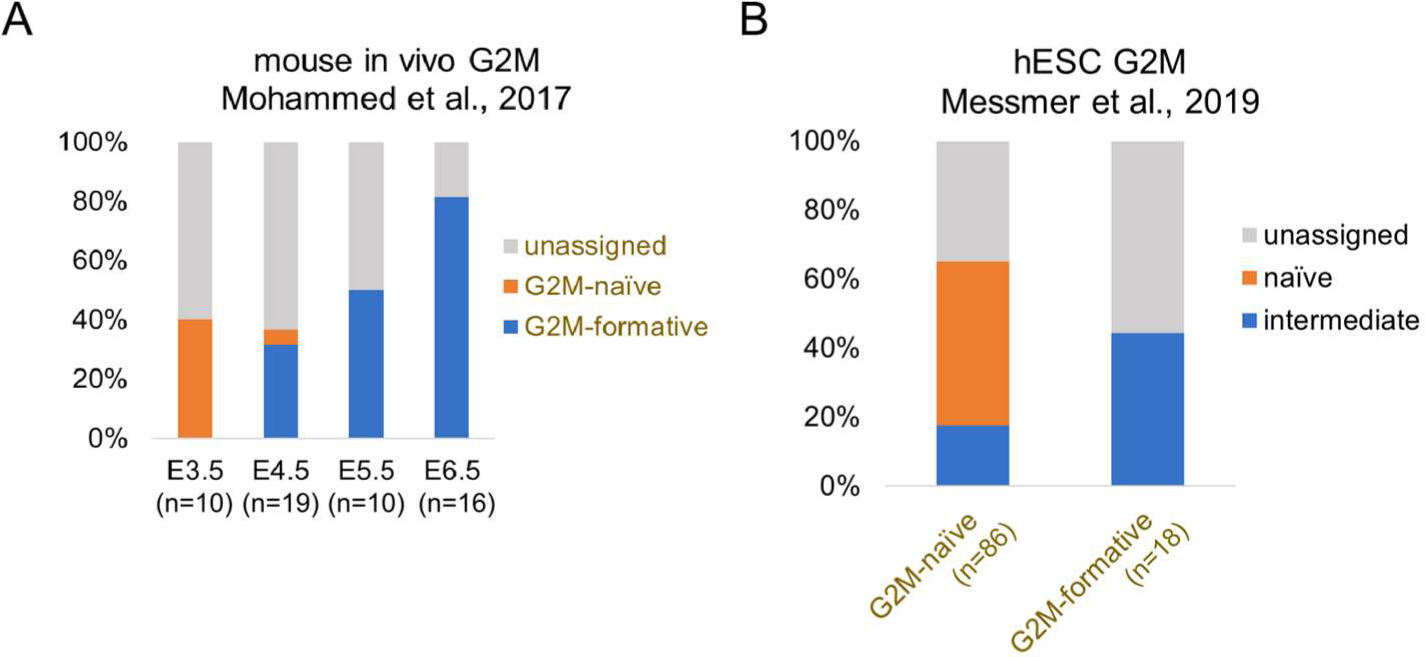
G2M sub-states separate the earliest pre-implantation stages. **A.** G2M-classified cells from the mouse embryo at distinct developmental stages (data from Mohammed et al., 2017) mapped to our **i**dentified G2M sub-states. More cells from the pre-implantation embryo (E3.5 and E4.5) map to the naïve state, while cells from the peri- and post-implantation embryos (E5.5 and E6.5) map to the formative state. **B.** Reverse mapping of our G2M sub-states to G2M-classified human ESCs (data from Messmer et al., 2019). Note the resemblance of the G2M-naïve mESCs with the G2M-naïve hESCs.

## DISCUSSION

### Developmental heterogeneity in G2M

It is well established that ESC developmental capacity is coordinated by cell cycle dynamics [8–11]. Accordingly, several single cell approaches have revealed pluripotent TF heterogeneity to partly result from differential cell cycle position [6,7,24,38]. In our study, we provide evidence for the temporal connection between cell cycle and TF heterogeneities by demonstrating developmental split of ESCs into distinct pluripotency sub-states specifically in G2M.

Heterogeneous expression of several pluripotency TFs in mESCs cultured in serum/LIF [2–5] has been proposed to reflect a variable developmental progression towards a primed state [39]. We show G2M cell cycle phase to separate mESCs grown in serum/LIF into TF expression states with reciprocal expression of naïve and early post-implantation factors, revealing developmental split of ESCs at later stages of the cell cycle. We did not see such strong bifurcation in G1 phase, which has been shown to be the phase when irreversible differentiation decisions are made [10]. The role of G2M in facilitating enhanced maintenance of pluripotency [8] as well as the earliest selection of ESC fate direction [9,40] corroborate our findings about the emergence of alternative pluripotency states in G2M.

We found that the most significant determinant of the mESC split into naïve versus formative state in G2M is the differential expression of TF Esrrb. Esrrb has been shown to be critical for the sustained activity of the naïve pluripotency regulatory network and in enhanced reprogramming to the naïve state [41]. Moreover, drop in Esrrb concentration below certain levels has been shown to trigger exit from naïve pluripotency [42]. Our finding that G2M cells exist either as Esrrb-H or Esrrb-L populations therefore strongly suggest G2M to be the phase when the decision to maintain or exit the naïve state is made.

Recently, S/G2M-controlled early ESC differentiation processes have received attention. mESC differentiation was shown to be initiated directly after DNA synthesis phase via mechanisms that involve delayed accumulation of repressive histone marks [11]. In hESCs, G2 cell cycle pause was required for initiation of differentiation [9]. A related phenomenon may be the enhanced capacity of G2M nuclei to reprogram cell fate via nuclear transfer [43], and higher reprogramming potency via cell fusion of ESCs in SG2 phase [44]. Therefore, G2M phase may facilitate change in cell fate in both directions, towards a more differentiated as well as less differentiated state.

Our most prominent finding that mESC exist in a significantly distinct pluripotency state with formative features in G2M builds a connection of ESC state control with the cell cycle-regulated chromatin dynamics. The formative state is hypothesised to be required for transcriptional and epigenetic remodelling to enable exit from the naïve state [1,30]. Remarkably, extensive ESC chromatin reorganisation occurs like a clockwork during each cell division. We hypothesise that the formative state in G2M may be linked to enhancer landscape switching to enable lineage specification [45,46]. This state diversion in G2M coincides with reduced Esrrb and other naïve TF expression, probably resulting in the decommissioning of the naïve enhancer landscape. This speculation is supported by findings about the role of Esrrb in activation of silent enhancers during a reverse developmental, the reprogramming process [47]. Taken together, we propose that the pluripotency state diversion in G2M may be mechanistically related to the extensive chromatin remodelling taking place in preparation for cell division that directly creates the epigenetic context for fate choices.

### G2M sub-states in vivo and between species

We found our G2M-naïve mESCs to match relatively well with the G2M cells from the pre-implantation mouse embryo, suggesting transcriptional recapitulation of the earliest stage of in vivo naivety.

Our preliminary comparison of G2M-phased mESCs and hESCs revealed noticeable similarity between these species. Transcriptomic profiling of the human embryo has demonstrated the expression of Esrrb and Klf2 only in the early inner cell mass which become downregulated in the pre-implantation epiblast, suggesting species-specific differences [48–50]. Nonetheless, in addition to the core pluripotency factors (Pou5f1, Sox2), the expression of the cell cycle regulators (Mybl2), chromatin remodelling factors (Jarid2) and factors promoting cell proliferation (Etv4/5) were shown to be conserved between rodents and human [48]. This may reflect lower sensitivity of G2M cells from different species to growth factor signalling and/or indicate partially redundant G2M-centred mechanisms of pluripotency maintenance.

### Conclusion

Single cell profiling of mESC expression states during cell cycle revealed the split of mESCs into naïve and formative pluripotency states in G2M. These findings demonstrate that mESC heterogeneity evident in serum/LIF conditions marks distinct developmental capacity attributable to transitions through late cell cycle phases.

## Author Contributions and Notes

K.J., I.C. and T.C. designed research; K.J., D.S., M.K., and A.S. performed research; K.J., D.S. and T.C. analyzed data; K.J. wrote the paper with contributions from D.S., M.K., T.C. and I.C. The authors declare no conflict of interest. This article contains supplementary information online:.

Supplementary_Figures.pdf

SupplTable1_SC3_markergenes.xls

SupplTable2_SCDE_z_scores.xlsx

## Acknowledgments

We thank Steve Pollard for help with gene editing experiment. This research was supported by Medical Research Council of the UK. K.J. was supported by the Mobilitas Pluss Programme grant (MOBTP28) in Estonia from the European Regional Development Fund. T.C. and D.S. were supported by a Chancellor’s Fellowship from the University of Edinburgh. D.S. was supported by MRC Doctoral Training Programme in Precision Medicine (#1805075).

