## Supplementary Figures for "G2M splits mouse embryonic stem cells into naïve and formative pluripotency states"

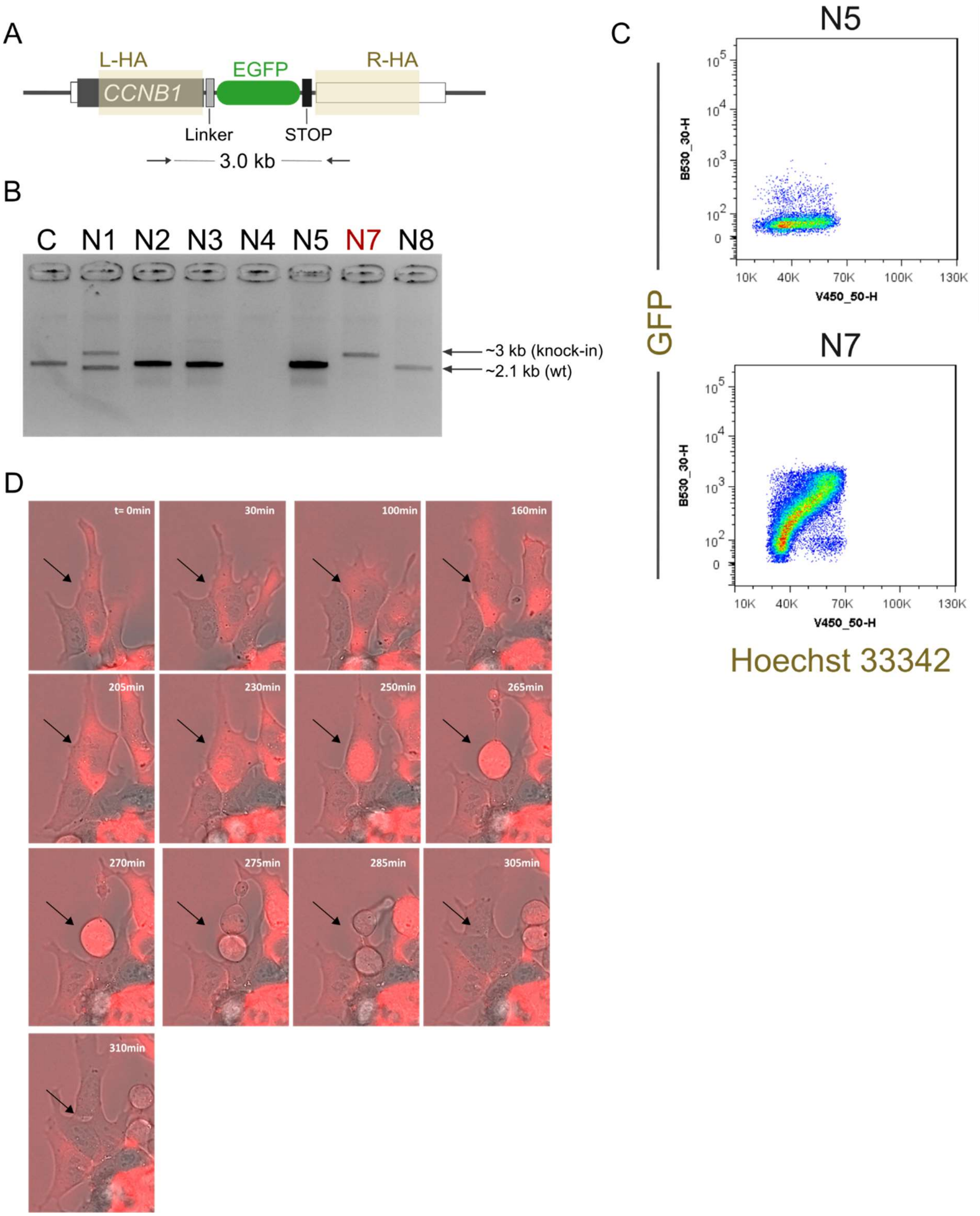


**Fig. S1. Generation of Cyclin B1-GFP reporter N7-mESC line. A.** A scheme of the *CCNB1* targeted locus where Cyclin B1-GFP fusion protein expression is driven by endogeneous *CCNB1* promoter. Left (L-HA) and right homology arms (R-HA) are highlighted. **B.** PCR-based genotyping of control (C) and selected targeted clones (N1-N8). Heterozygous (N1), homozygous (N7) and non-targeted (N2, N3, N5, N8) clones were recovered. **C.** Flow cytometry dot-blot confirming near-linear correlation between CyclinB1-GFP expression and DNA content (by live Hoechst 33342 staining) only in targeted (N7) mESCs. **D.** Time-lapse images of N7-mESCs showing nuclear accumulation of CyclinB1:GFP in mitosis (t = 250-270 min) and rapid loss of expression immediately after cell division (t = 275 min). Black arrow indicates the representative cell going through cell division.

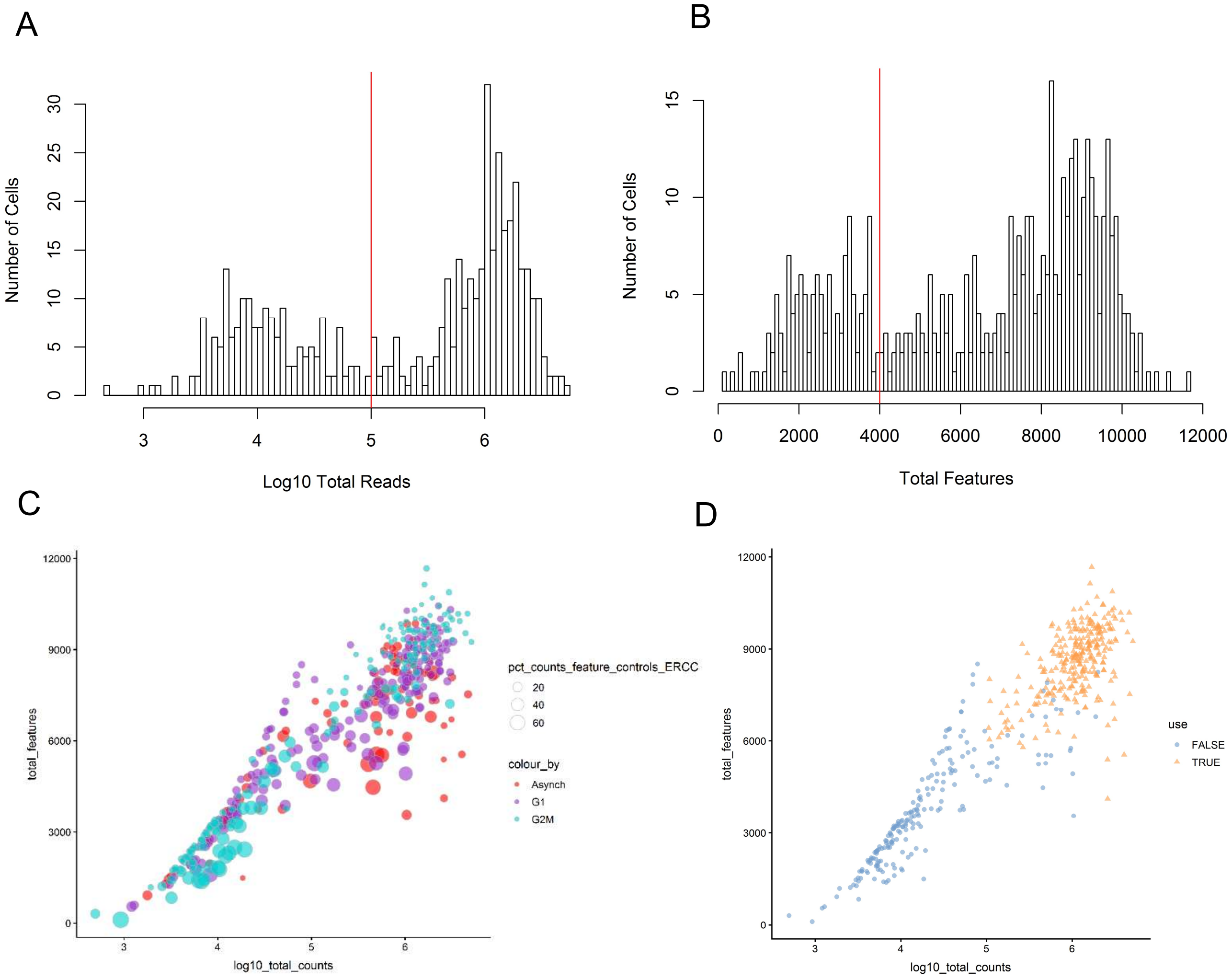


**Fig. S2. Quality filtering of single-cell data A.** Histogram of log10 total reads for asynchronous, G1 and G2M mESCs. Red line denotes quality control (QC) cut off, where cells with less than $10^5$ reads were rejected. **B.** Histogram of total features (genes) for asynchronous, G1 and G2M mESCs. Red line denotes QC cut off, where cells with less than 4000 features were rejected. **C and D.** Scatter plots showing Log10 total counts (reads per cell) on x-axis against total features (genes per cell) on y-axis for asynchronous, G1 and G2M mESCs. Cell points in A vary in size based on percentage of reads that contain ERCCs and are coloured by FACS-sorted population. In B, cells are coloured by which have passed (TRUE) or failed (FALSE) QC measures.

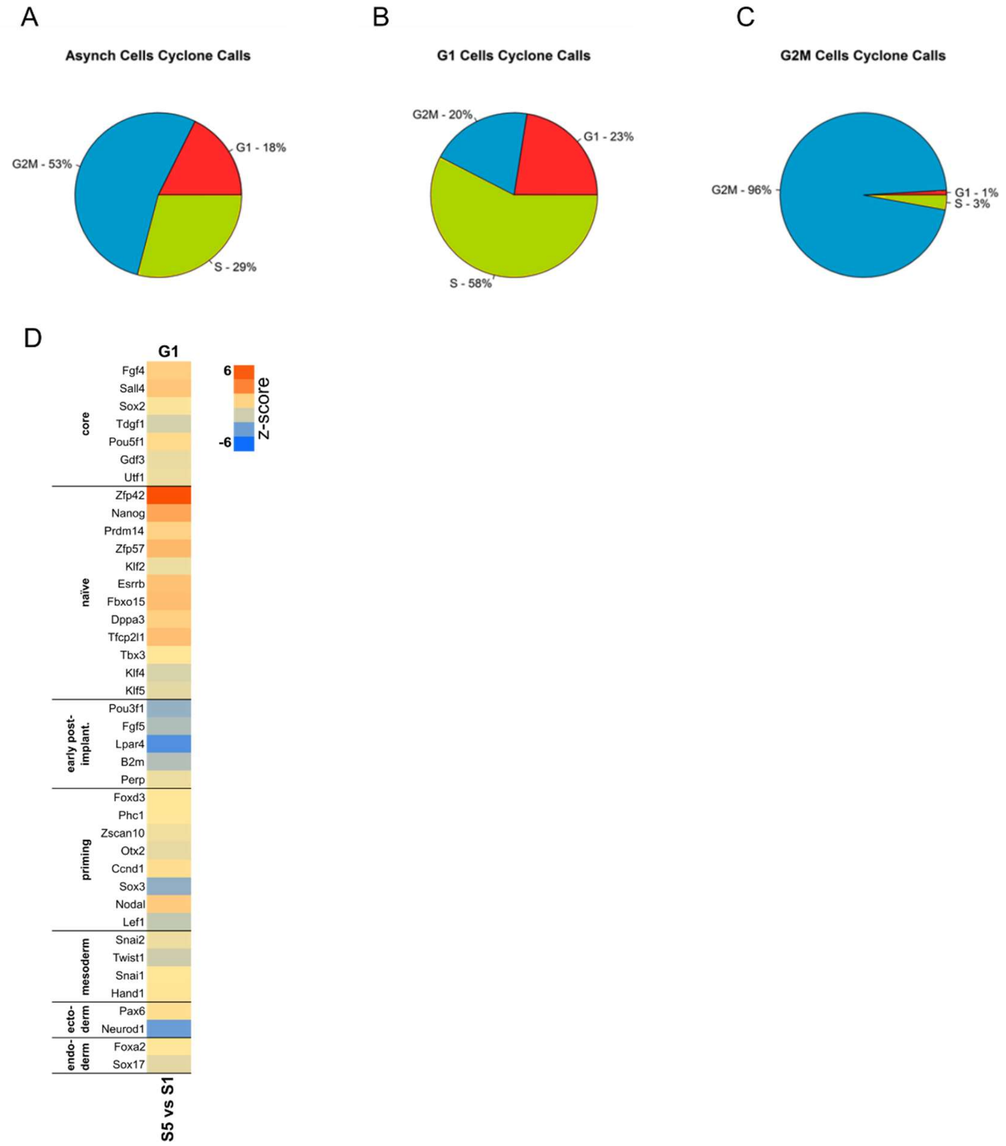


**Fig. S3. Cell cycle phase quantification of sorted cells.** The distribution of cells in **A.** asynchronous **B.** G1-sorted and **C.** G2M-sorted population into Cyclone-assigned cell cycle phases showing high purity of G2M cells. **D.** SCDE analysis (z-scores) between G1 sub-states showing less pronounced reverse expression between naïve and early post-implantation genes (except Zfp42 and Lpar4).
